# Synchronized proinsulin trafficking reveals delayed Golgi export accompanies β-cell secretory dysfunction in a rodent model of hyperglycemia

**DOI:** 10.1101/2022.10.31.514578

**Authors:** Cierra K. Boyer, Casey J. Bauchle, Jianchao Zhang, Yanzhuang Wang, Samuel B. Stephens

## Abstract

The pancreatic islet β-cell’s preference for release of newly synthesized insulin requires careful coordination of insulin exocytosis with sufficient insulin granule production to ensure that insulin stores exceed peripheral demands for glucose homeostasis. Thus, the cellular mechanisms regulating insulin granule production are critical to maintaining β-cell function. In this report, we utilized the synchronous protein trafficking system, RUSH, in primary β-cells to evaluate proinsulin transit through the secretory pathway leading to insulin granule formation. We demonstrate that the trafficking, processing, and secretion of the proinsulin RUSH reporter, proCpepRUSH, are consistent with current models of insulin maturation and release. Using a rodent dietary model of hyperglycemia and β-cell dysfunction, we show that proinsulin trafficking is impeded at the Golgi and coincides with the decreased appearance of nascent insulin granules at the plasma membrane. Ultrastructural analysis of β-cells from diabetic leptin receptor deficient mice revealed gross morphological changes in Golgi structure, including shortened and swollen cisternae, and partial Golgi vesiculation, which are consistent with defects in secretory protein export. Collectively, this work highlights the utility of the proCpepRUSH reporter in studying proinsulin trafficking dynamics and suggests that altered Golgi export function contributes to β-cell secretory defects in the pathogenesis of Type 2 diabetes.

## Introduction

Pancreatic islet β-cells regulate glucose homeostasis through controlled release of the hormone, insulin, to promote nutrient uptake and storage in peripheral tissues, such as skeletal muscle, liver, and adipose. In the β-cell, insulin is stored in dense core secretory granules, which undergo regulated exocytosis in response to metabolic cycles that drive Ca^2+^-dependent fusion of the insulin granule with the plasma membrane^1,2^. A β-cell contains 8,000-10,000 insulin secretory granules at steady state and can secrete 5-10 % of the insulin content per hour in response to nutrient stimulation^3,4^. β-cells preferentially secrete (∼60%) newly synthesized insulin (formed within 24 h), despite the extended half-life (∼2.7 days) of an insulin granule^5,6^; however, the molecular determinants for this preference are not well understood.

To maintain the steady supply of releasable insulin granules, the β-cell devotes 30-50% of its total protein biosynthetic capacity to insulin production^7–9^. From the initial synthesis of proinsulin, to the endoproteolytic conversion to insulin in the maturing granule, pulse-chase radiolabeling studies estimate insulin biosynthesis requires approximately 3 hours^5^. Proinsulin folding in the endoplasmic reticulum (ER) is widely regarded as a rate-limiting step in the secretory biosynthetic pathway and recent data highlights that impairments in proinsulin folding and disulfide bond formation limit insulin granule formation in diabetes models^10–12^. In addition, proinsulin exit from the *trans*-Golgi network (TGN) may also be a key regulated step in the insulin biosynthetic pathway^13,14^. While proteins, such as proinsulin, lack well-defined topological signals for vesicle sorting, protein cargo entering the regulated secretory pathway may be actively sorted in the TGN prior to delivery into the budding granule^15–17^. Indeed, granin proteins, VGF and chromogranin B (CgB), are required for efficient proinsulin exit from the TGN^18,19^ and may facilitate pH-dependent proinsulin condensation within the Golgi cisternae as a key step in proinsulin sorting^20^. Despite these advances, molecular determinants of proinsulin transit through the secretory system are only recently being deciphered^21,22^ and the specificity of these interactions for proinsulin versus other maturing secretory cargo has not been investigated. This is particularly important as both major forms of diabetes, Type 1 and 2 (T1D and T2D), are known to present with defects in proinsulin processing and insulin storage that may reflect alterations early in the proinsulin trafficking pathway that bypass critical checkpoints for proinsulin packaging^8,10,23–25^.

Historically, studies of proinsulin trafficking have been challenging because techniques to temporally resolve critical steps in proinsulin maturation and granule formation *in situ* were reliant on conventional radioisotope pulse-chase labeling and electron microscopy^14,26^. With the development of fluorescence-based trafficking reporters such as SNAPtag, pulse-chase labeling has revealed age-dependent alterations in granule mobility^27^ and recently demonstrated delayed exit of proinsulin from the ER in rodent models of hyperglycemia and diet-induced pre-diabetes^12^. In this report, we utilized the *in situ* fluorescent pulse-chase approach termed Retention Using Selective Hooks (RUSH)^28^ to study proinsulin trafficking and insulin granule formation in primary β-cells. The RUSH system allows for synchronous monitoring of protein trafficking through the secretory pathway and is based on the reversible retention of a reporter protein (proinsulin) fused to streptavidin-binding peptide (SBP), which strongly interacts with an organelle-localized hook containing streptavidin (SA)^28^. Trafficking can be initiated by biotin addition to the culture media, which rapidly diffuses into cellular compartments, outcompetes SBP binding to SA, and thereby releases the cargo reporter to follow its natural process through the secretory pathway. Past studies using RUSH reporters to examine TGN exit have relied on surrogate markers, such as secretogranin II and neuropeptide Y (NPY), to investigate soluble proteins trafficking dynamics in insulinoma cells^29^; however, our study provides the first example of this technique with proinsulin in primary β-cells. Here, we show the trafficking of a proinsulin RUSH reporter, termed proCpepRUSH, from the ER to the secretory granule occurs within 2 h. These data are consistent with the estimated 3 h timeline for completed endoproteolytic conversion of proinsulin to insulin, which is known to occur within the maturing secretory granule^14^. We further demonstrate the processing of proCpepRUSH occurs post-Golgi exit and that only fully processed, CpepRUSH, is secreted from β-cells in response to glucose stimulation. Using a rodent dietary model of β-cell dysfunction, we show that proinsulin trafficking is altered in primary β-cells, with a specific delay in Golgi export. This delay coincides with substantial alterations to Golgi structure observed in leptin-receptor deficient db/db mice. Ultrastructural analysis of Golgi stacks revealed shortened, dilated cisternae and a high degree of Golgi vesiculation. Collectively, this work demonstrates the value of the proCpepRUSH reporter in studying proinsulin trafficking and suggest that alterations in Golgi export function contribute to granule trafficking delays during the pathogenesis of β-cell dysfunction in T2D.

## Results

### Synchronized trafficking of proinsulin

In this study, we developed a proinsulin trafficking reporter based on the Retention Using Selective Hooks (RUSH) design^28^. We used ER-targeted streptavidin, by virtue of a C-terminal KDEL sequence (SA-KDEL), as the organelle localized hook to synchronize proinsulin release from the ER and monitor proinsulin trafficking through the Golgi and into the mature insulin secretory granule (Fig. 1A). To generate the proinsulin reporter, we inserted streptavidin-binding peptide (SBP) and superfolder GFP (sfGFP) within the C-peptide region of human preproinsulin (proCpepRUSH). This region avoids potential proinsulin folding and aggregation problems^30^ and has been successfully used for other proinsulin reporters^13,19,31,32^. For β-cell specific expression in primary islets, we generated a recombinant adenovirus using the rat insulin promoter (RIP) driving expression of a bicistronic cassette, containing SA-KDEL, followed by the EMCV IRES and proCpepRUSH. This approach guarantees that both constructs (SA-KDEL and proCpepRUSH) are present in the same β-cell and that SA-KDEL is in molar excess to proCpepRUSH, which ensures high fidelity of the trafficking system.

**Figure 1.**
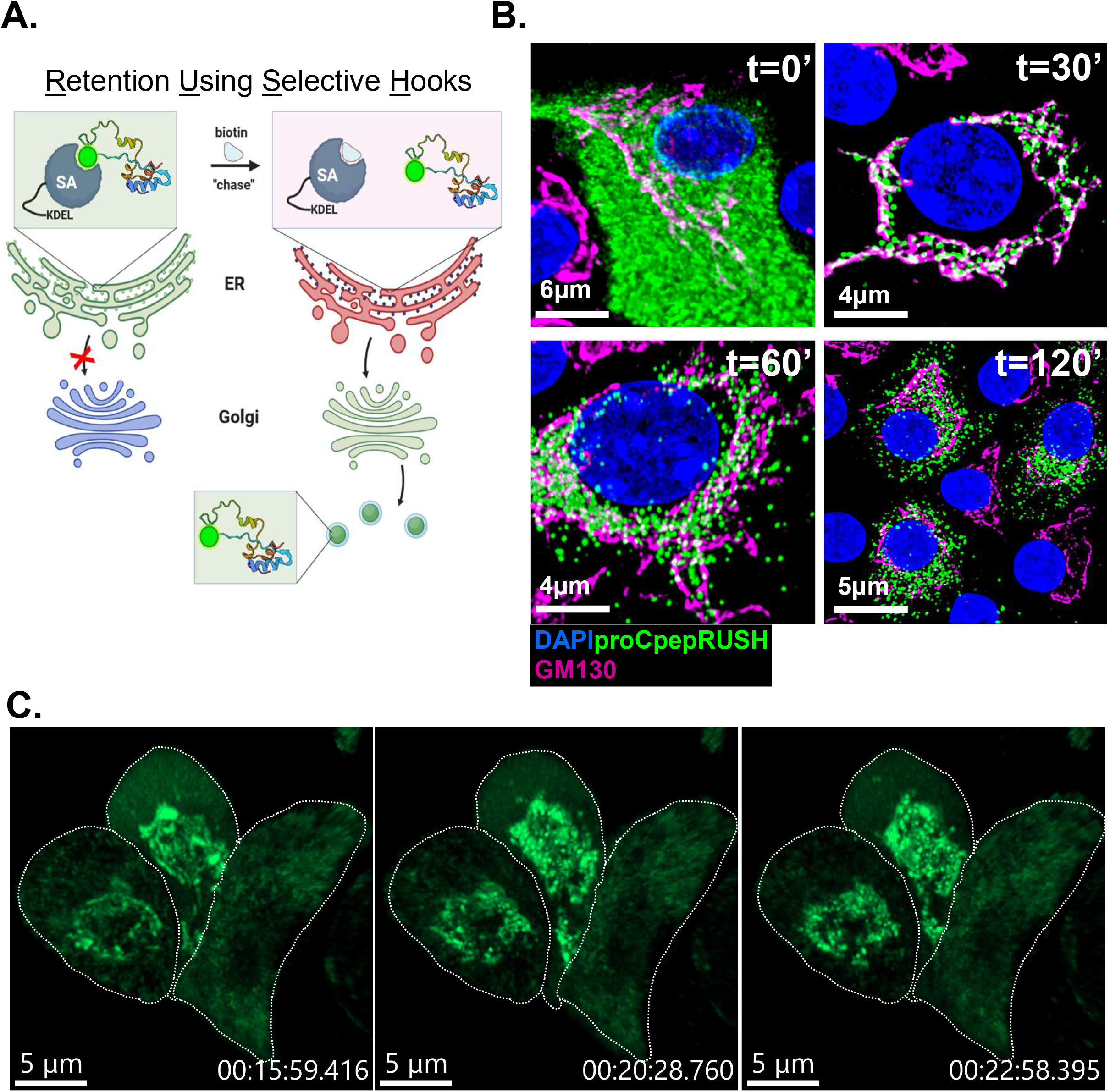
Proinsulin trafficking using RUSH in primary mouse β-cells. (**A**) Schematic of proCpepRUSH trafficking was created using biorender.com. (**B**) Mouse islets (C57BL6/J) treated with AdRIP-proCpepRUSH were examined 48 h post-infection. Biotin addition (200 μM) was used to initiate proCpepRUSH (green) trafficking. Islet cells were fixed at the indicated times post-biotin addition and immunostained for GM130 (magenta) and counterstained with DAPI (blue). (**C**) Specific time points from time-lapse imaging (Movie S1) of proCpepRUSH trafficking are shown with cell outlines.

Using the RUSH system (SA-KDEL and proCpepRUSH), synchronous release of proinsulin was examined in primary mouse islets. Prior to biotin administration, proCpepRUSH was present throughout the lacy network of the cell body (Fig. 1B, t=0) and precisely mirrors SA-KDEL immunostaining (Fig. S1A). This staining pattern is consistent with ER localization and likely reflects the tight association of proCpepRUSH-containing SBP and the ER-localized SA-KDEL. Following biotin addition, proCpepRUSH was observed in the Golgi region (indicated by GM130, magenta) within 30 min, and remained proximal to the Golgi for an additional 30 min (t = 60min) (Fig. 1B, Movie S1). At 120 min post-biotin addition, numerous proCpepRUSH puncta were evident throughout the cell body distinct from the Golgi (Fig 1B) and the ER (Fig. S1B) suggesting that proCpepRUSH-containing vesicles had been released from the Golgi and entered the pool of insulin storage granules (Movie S1). Upon further examination of Golgi transit, we noted the appearance of proinsulin in the Golgi structure was initially diffuse (Fig 1C, t=15 min and Movie S1), but transitioned to discrete regions of high-density staining (Fig. 1B, t=30 min and Fig.1C, t=20 and 23 min). These data are consistent with studies demonstrating that proinsulin condenses in the Golgi prior to delivery into the budding secretory granule^14,20^. To confirm this finding, we used a pan-insulin antibody that detects proinsulin and observed discrete regions of high-density proinsulin staining within the Golgi (GM130) volume in primary mouse β-cells (Fig. S1C-F). Collectively, our timeline for proCpepRUSH trafficking from the ER to the granule is consistent with the known kinetics of proinsulin conversion to mature insulin^5,33^.

### Processing and secretion of proCpepRUSH

To verify the successful trafficking of the proCpepRUSH system, we used immunostaining to examine the localization in INS1 832/3 insulinoma cells pre- and post-biotin addition. In the absence of biotin treatment (t=0), proCpepRUSH was diffusely present throughout the cell body (Fig. 2A), similar to primary β-cells (Fig. 1B, t=0). We observed strong co-localization of proCpepRUSH with the ER chaperone, GRP94 (red) that was distinct from the Golgi marker, TGN38 (magenta) (Fig. 2A). Post-biotin treatment (t=180 min), proCpepRUSH puncta were present in carboxypeptidase E (CPE)-positive vesicles (red), consistent with successful trafficking to insulin secretory granules (Fig. 2B). Note, that we would not expect all CPE-positive vesicles to contain proCpepRUSH because many of the CPE-positive vesicles would have been formed prior to biotin-stimulated trafficking of our proCpepRUSH construct. In contrast to CPE, proCpepRUSH was clearly absent from endolysosomal vesicles containing the mannose-6-phosphate receptor (M6PR, red) (Fig. 2C), suggesting that proCpepRUSH was correctly targeted to the regulated secretory pathway.

**Figure 2.**
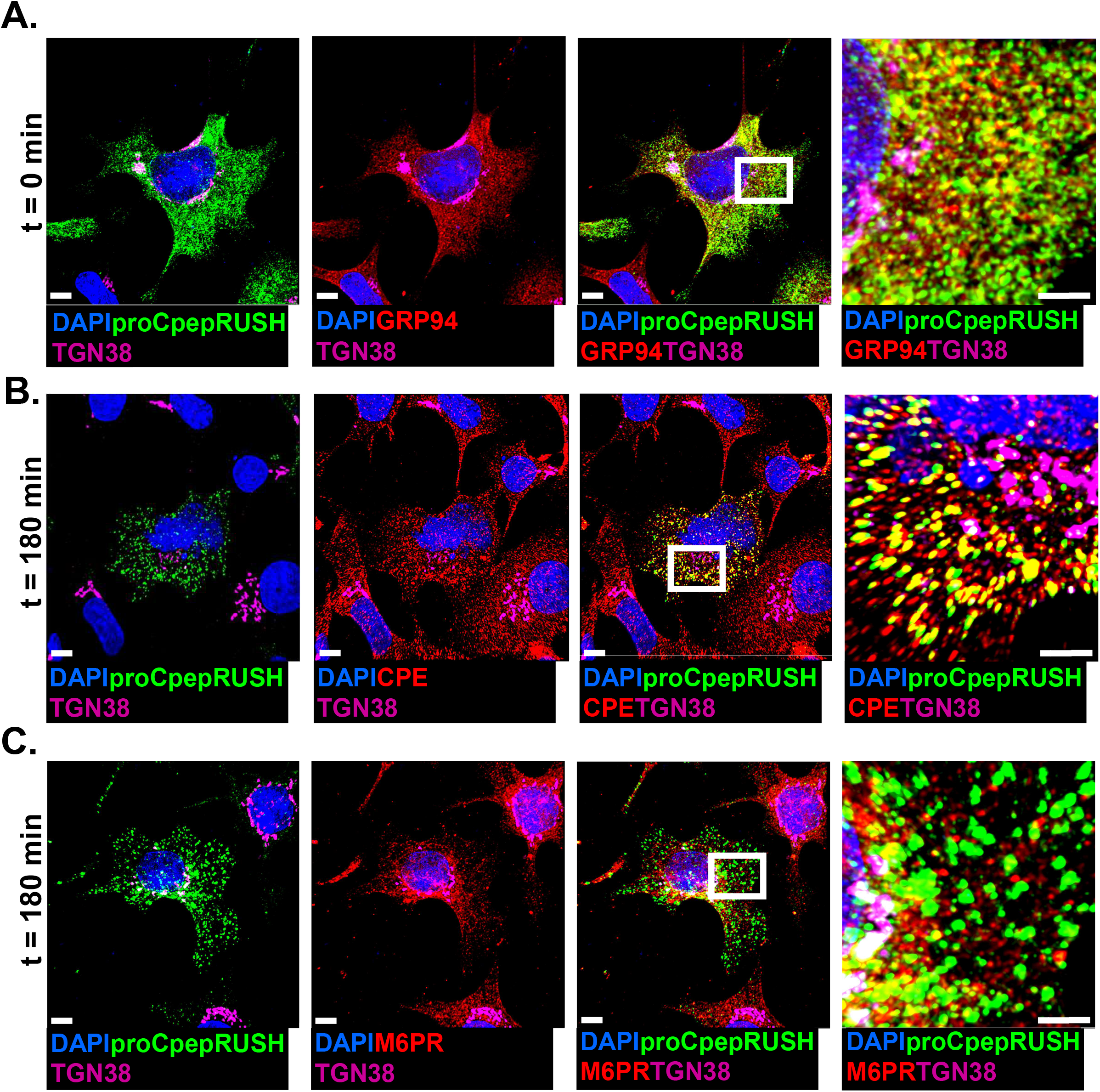
Co-localization marker analysis of proCpepRUSH trafficking. INS1 832/3 cells expressing AdRIP-proCpepRUSH were treated with biotin (200 μM) to initiate proCpepRUSH (green) trafficking for the indicated times prior to fixation. Cells were immunostained for GRP94 (red) (**A**), mannose-6-phosphate receptor (M6PR, red) (**B**) or CPE (red) (**C**). (**A**-**C**) TGN38 (magenta) and DAPI (blue) were used as counterstains. Boxed insets are shown with additional magnification. Scale bar = 5 μm.

To determine if proCpepRUSH was available for regulated secretion, INS1 832/3 cells expressing proCpepRUSH were treated with biotin for 3 h to stimulate proCpepRUSH trafficking into secretory granules, followed by culture at either basal (2.5 mM) or stimulatory (12 mM) glucose for an additional 3 h to elicit insulin and proCpepRUSH exocytosis. Using immunoblot analysis, fully processed CpepRUSH (predicted molecular weight is 34.3 kDa), was identified in the media from glucose-stimulated (12 mM) cells, and not present in cell media following basal glucose culture or no virus controls (Fig. 3A). Note that both fully processed CpepRUSH and non-processed proCpepRUSH (predicted molecular weight is 40.7 kDa), were present in cell lysates, but only processed CpepRUSH was secreted. Importantly, proCpepRUSH expression does not interfere with normal β-cell function (Fig. 3B).

**Figure 3.**
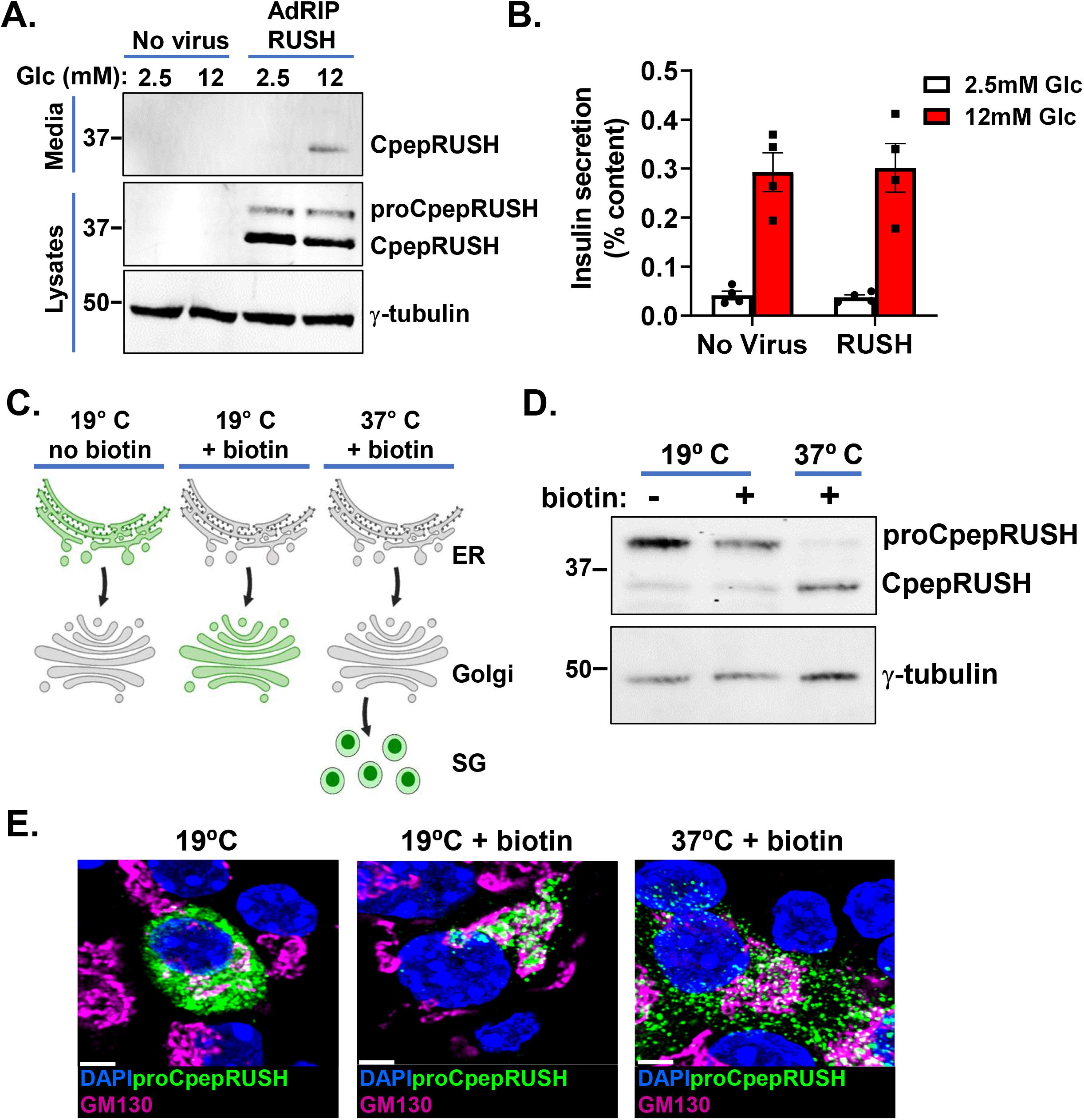
Processing and secretion of proCpepRUSH. (**A**) INS1 832/3 cells treated with AdRIP-proCpepRUSH or no virus (control) were supplemented with biotin (200 μM) for 3 h Glucose-stimulated insulin secretion was measured by static incubation in media containing 2.5 mM Glc or 12 mM Glc for 3 h each. TCA precipitated media was analyzed by immunoblot and compared to whole cell lysate. (**B**) Primary mouse islets (C57BL6/J) treated with AdRIP-proCpepRUSH or no virus were analyzed for glucose-stimulated insulin secretion via sequential static incubation in media containing 2.5 mM and 12 mM glucose for 1 h each as indicated. Data are normalized to insulin content determined from cell lysates. Data represents the mean ± S.E.M. (**C**) Schematic demonstrating the use of biotin treatment in conjunction with temperature block to examine ER, Golgi, and secretory granule localization of proCpepRUSH. This model was created using biorender.com. (**D, E**) Isolated mouse islets were treated with AdRIP-proCpepRUSH. 48 h post-infection, islets were incubated at 19° C for 1 h (ER retained), then treated with biotin (200 μM) for an additional 2 h at 19° C to initiate proCpepRUSH (green) trafficking, but prevent Golgi exit. Islets were then shifted to 37° C for 2 h to stimulate Golgi release. (**D**) Whole cell lysates were analyzed by immunoblot using anti-GFP to detect proCpepRUSH and CpepRUSH. (**E**) Cells were immunostained for GM130 (magenta) and counterstained with DAPI (blue). Scale bar = 5 μm.

Furthermore, similar to another proinsulin reporter, proCpepSNAP, expression of proCpepRUSH does not elicit an ER stress response as compared to the SERCA2 inhibitor, thapsigargin, which strongly increases expression of UPR targets, BiP, CHOP, GADD34, and XBP-1 (s/u) (Fig. S2). To demonstrate successful processing of proCpepRUSH occurred in the maturing granule and not earlier in the secretory pathway, we separately evaluated proCpepRUSH in the ER, Golgi, and secretory granule. For these studies, we used biotin treatment to initiate proCpepRUSH trafficking from the ER to Golgi in conjunction with a temperature block (19° C) to prevent TGN exit (Fig. 3C)^34,35^. Under conditions where proCpepRUSH staining resided primarily in the ER (19° C) or Golgi (19° C with biotin) (Fig. 3C, E), proCpepRUSH was identified as the full-length prohormone (Fig. 3D). Following Golgi release by shifting cells to 37° C, proCpepRUSH was trafficked into secretory granules (Fig. 3C, E) and proteolytically processed to CpepRUSH (Fig. 3D). Note that in approximately 5-10% of the cells, proCpepRUSH was observed as granular (rather than ER) prior to biotin addition (data not shown), which is consistent with immunoblot detection of processed CpepRUSH in the non-biotin treated samples (Fig. 3D). Collectively, these data demonstrate that the trafficking, processing, and secretion of proCpepRUSH/ CpepRUSH are consistent with the regulation of endogenous proinsulin/ insulin.

### Impaired proinsulin trafficking occurs in a rodent model of β-cell dysfunction

To demonstrate the utility of the proCpepRUSH reporter, we examined proinsulin trafficking in a rodent dietary model of early-stage diabetes. Here, we used primary islets from male C57BL6/J mice placed on a Western diet (WD; 40% fat/kcal, 43% carbohydrate/kcal) or standard chow (SC). By 8 weeks, increased body weight (Fig. 4A) and ad lib fed hyperglycemia (Fig. 4B) were evident in mice on WD compared to SC controls. In addition, reduced glucose tolerance (Fig. 4C), fasting hyperinsulinemia (Fig. 4D), and impaired glucose-stimulated insulin release (Fig. 4D) were observed in WD fed mice. These data are consistent with defects in β-cell function occurring early in the development of diabetes ^36,37^.

**Figure 4.**
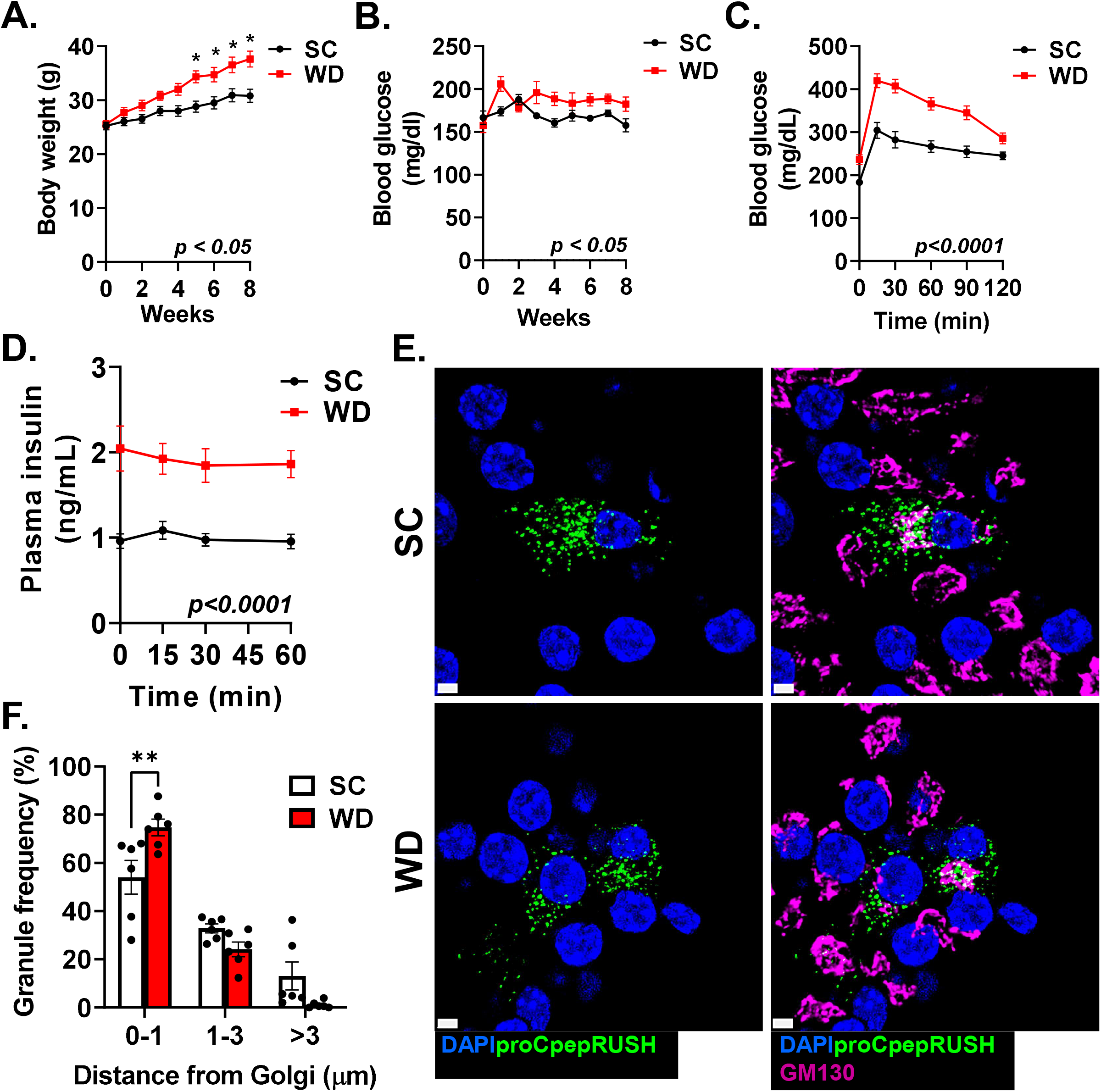
Impaired insulin granule formation in a diet-induced model of islet dysfunction. 8-10 week old male C57BL6/J mice were maintained on standard chow (SC) or Western diet (WD; 40% fat/kcal, 43% carbohydrate) for 8 weeks. Body weight (**A**) and ad lib fed blood glucose (**B**) were measured weekly (n=10 per group). (**C-D**) Following 8 weeks of diet, 4-6 h fasted mice (n=14-15 per group) were injected i.p. with glucose (1 mg/g bw). Blood glucose (**C**) and plasma insulin (**D**) were measured at the indicated times. (**E-F**) Isolated mouse islets were treated with AdRIP-proCpepRUSH (n=6 mice per group). 48 h post-infection, islets were incubated with biotin (200 μM) for 3 h at 37° C to stimulate proCpepRUSH (green) trafficking. Cells were immunostained for GM130 (magenta) and counterstained with DAPI (blue). (**E**) Representative images are shown. Scale bar = 5 μm. (**F**) The distances of proCpepGFP-positive granules from the Golgi were normalized per cell as a frequency distribution. Data represents the mean ± S.E.M. *p < 0.05 by 2 way-ANOVA with repeated measures (**A**-**D**) or Sidak post-test analysis (**F**).

Using the proCpepRUSH system, we examined proinsulin trafficking 3 h post-biotin addition to determine if defects in insulin granule formation could be detected in β-cells from WD fed mice. We used proCpepRUSH granule distance from the Golgi to assess trafficking and normalized the data as a relative frequency of granule distances per cell imaged, which accounts for variations in reporter expression between cells and conditions^12,19^. As shown in Figure 4E, proCpepRUSH-positive granules (green) in β-cells from SC fed mice were distributed throughout the cell body. In contrast, proCpepRUSH-positive granules from WD fed mice were commonly clustered around the Golgi (GM130, magenta), consistent with our previous data^12^. Quantitation of the granule distances to the nearest point on the Golgi surface (GM130) revealed a significant increase in the frequency (75 %) of nascent proCpepRUSH-labeled granules less than 1 µm from the Golgi in β-cells from WD compared to SC fed mice and a corresponding trend toward decreasing granule numbers greater than 3 µm from the Golgi (Fig. 4F).

Our prior work using a separate proinsulin reporter, proCpepSNAP, demonstrated that insulin granule formation was impaired in diabetes models due to delayed ER-Golgi transit of proinsulin^12^. Whether an additional delay in proinsulin trafficking from the Golgi also occurs and contributes to the insulin granule deficit is not known. To address this, we took advantage of the temperature dependence for Golgi release^34,35^ that we previously used to examine proCpepRUSH processing (Fig. 3D, E). Primary islets from SC and WD fed mice (8-10 wks) were cultured at 19° C for 2 h in the presence of biotin to promote proCpepRUSH accumulation in the Golgi, but prevent Golgi exit (Fig. 5A; Golgi block -white arrows denote Golgi-localized proCpepRUSH). Following a 2 h shift to 37° C, we examined proinsulin release from the Golgi (Fig. 5A; Golgi release). We measured the distances of proCpepRUSH-positive granules from the nearest point on the Golgi (GM130) normalized per cell as a measure of granule trafficking (Fig. 5B). In β-cells from SC fed mice, nascent proCpepRUSH granules (green) were evident throughout the cell body with approximately 40 % of the nascent granules within 1 µm of the Golgi (GM130) and ∼20% of nascent granules greater than 3 µm from the Golgi. In contrast, β-cells from WD fed mice presented with ∼68 % of proCpepRUSH-positive granules proximal (< 1 µm) to the Golgi and fewer than 4 % beyond 3 µm from the Golgi. TIRF microscopy was used to corroborate these findings and revealed a greater than 50 % reduction in the presence of nascent proCpepRUSH-labeled granules localized to the plasma membrane (≤ 150 nm) in the WD model (Fig. 5C-D). Together, these data suggest that proinsulin trafficking from the Golgi is delayed in a rodent dietary model of β-cell dysfunction.

**Figure 5.**
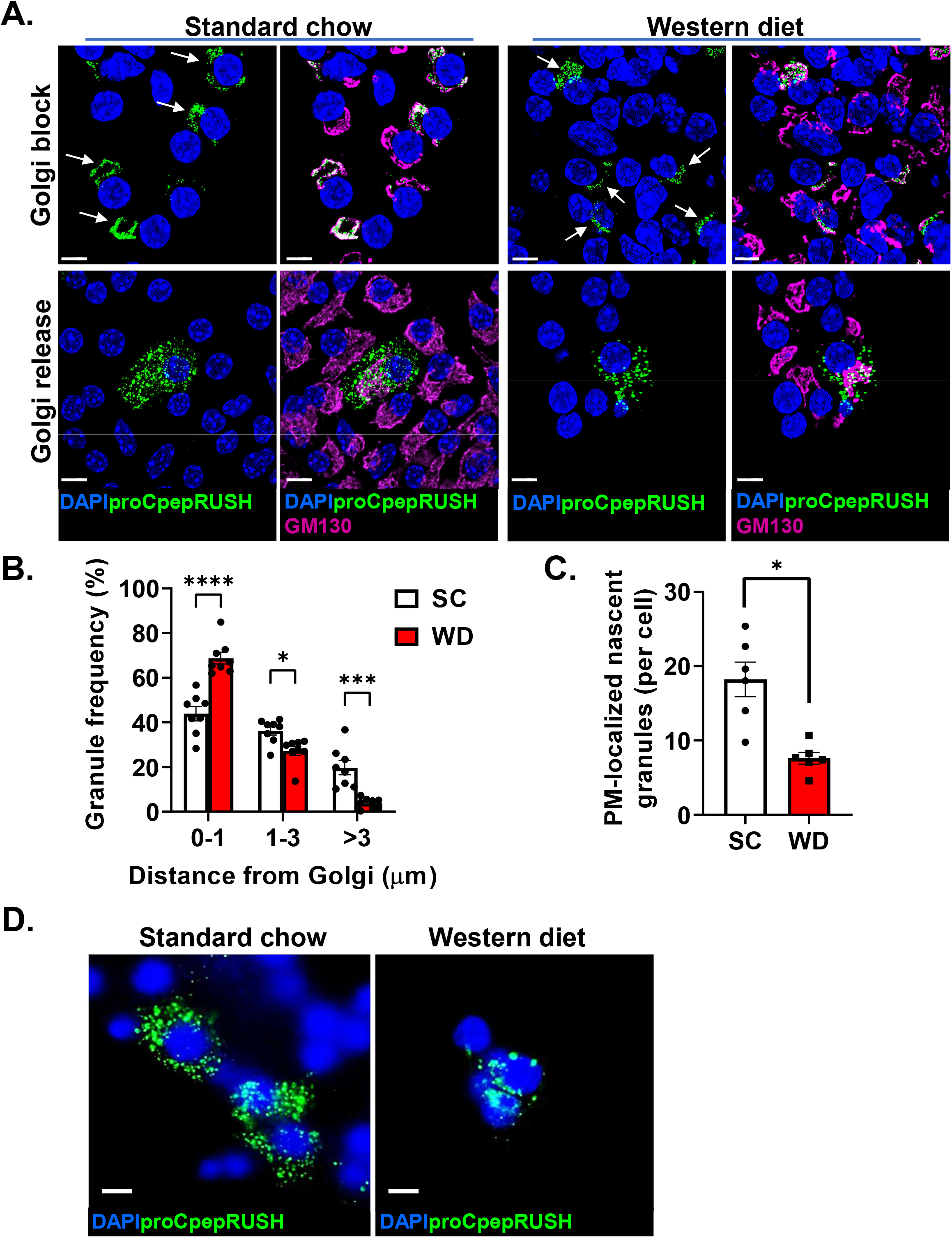
Impaired insulin trafficking from the Golgi in a diet-induced model of islet dysfunction. 8-10 week old male C57BL6/J mice were maintained on standard chow (SC) or Western diet (WD; 40% fat/kcal, 43% carbohydrate) for 8-10 weeks. Isolated mouse islets were treated with AdRIP-proCpepRUSH. 48 h post-infection, islets were incubated at 19° C for 1 h and treated with biotin (200 μM) for an additional 2 h at 19° C to initiate proCpepRUSH (green) trafficking, but block Golgi exit. Islets were then shifted to 37° C for 2 h to stimulate Golgi release. Cells were immunostained for GM130 (magenta) and counterstained with DAPI (blue). (**A, B**) Islets were imaged by confocal microscopy. (**A**) Representative images are shown for Golgi block (white arrows denote Golgi-localized proCpepRUSH) and Golgi release. (**B**) The distances of proCpepRUSH-positive granules from the Golgi (2 h post-Golgi release) were normalized as a frequency distribution per cell (n=8 mice per group). (**C, D**) Islets were imaged by TIRF microscopy following 2 h Golgi release. (**C**) The total number of plasma membrane (PM)-localized proCpepRUSH-positive granules were normalized per cell (n=6 mice per group). (**D**) Representative TIRF images are shown. (**B, C**) Data represents the mean ± S.E.M. *p < 0.05 by 2 way-ANOVA with Sidak post-test analysis (**B**) or two-tailed unpaired Student t test (**C**). (**A, D**) Scale bar = 5 μm.

Remodeling the Golgi cisternae is well-documented to directly impact secretory protein export. To explore this feature, we investigated Golgi morphology in primary β-cells from a genetic model of diabetes, the leptin receptor deficient mouse (*Lepr*^*db/db*^, db/db). By 14 wks of age, db/db mice displayed a significant impairment in glucose homeostasis (Fig. 6A), hyperinsulinemia (Fig. 6B), and decreased β-cell response to glucose challenge (Fig. 6B). Ultrastructural analysis of healthy β-cells from db/+ mice revealed thin Golgi cisternae that were organized into long stacks (Fig. 6C), with the cisternal length greater than 1 μm (Fig. 6D) and averaging 6 cisternae per stack (Fig. 6E). In contrast, Golgi structure in diabetic β-cells (db/db) was disorganized (Fig. 6C). Golgi cisternae were shorter in length (0.8 μm; Fig. 6D), presented with fewer cisternae per Golgi stack (average of 5; Fig. 6E) and more Golgi stacks per β-cell (Fig. 6F). In addition, we observed significant distention of the individual Golgi cisterna in db/db β-cells (Fig. 6G) and an overall increase in Golgi vesiculation (Fig. 6H). Collectively, these data are consistent with protein export defects and suggest that disruptions in Golgi structure may contribute to insulin trafficking defects in diabetes models.

**Figure 6.**
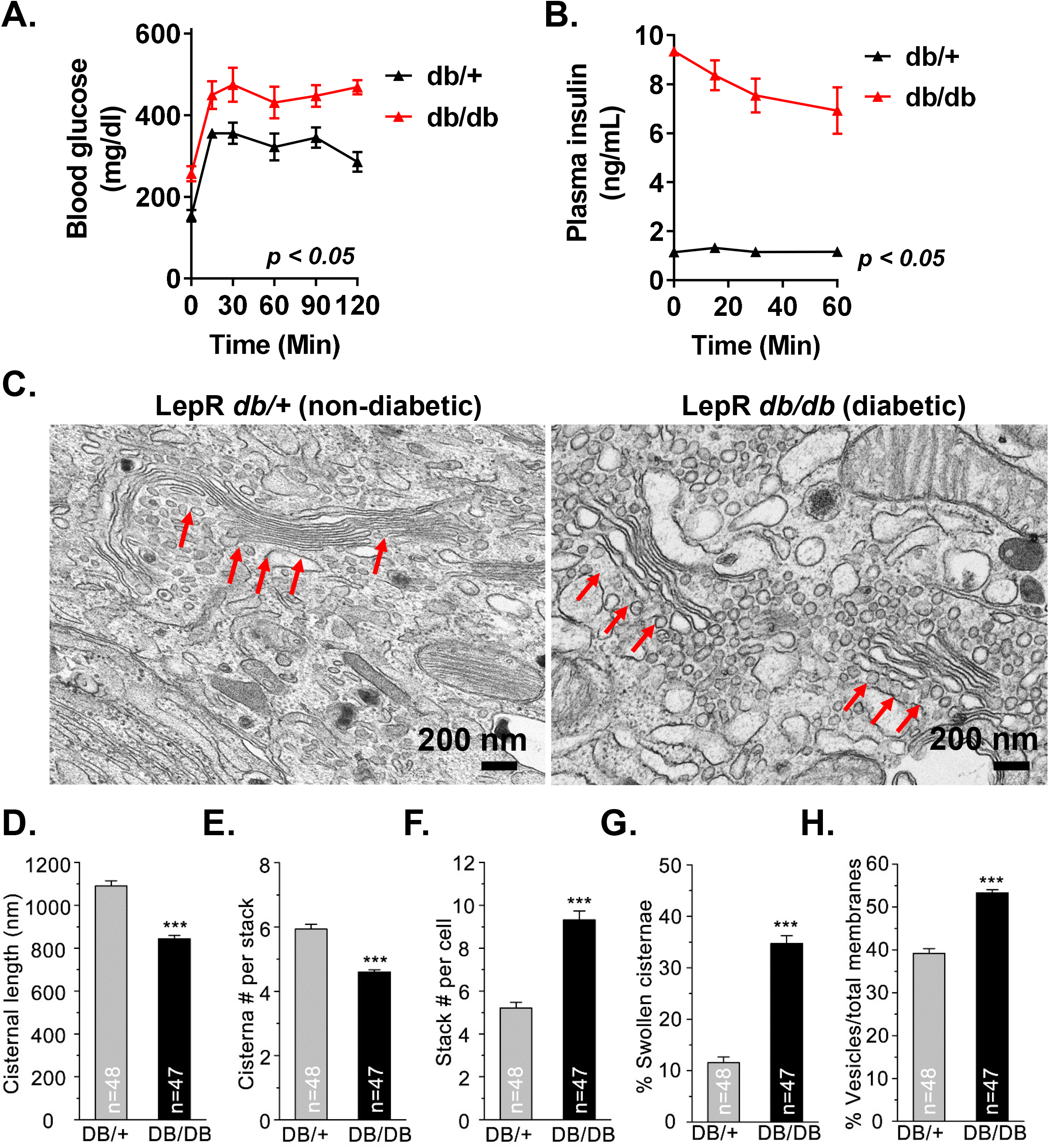
Alterations in Golgi morphology in a rodent model of islet dysfunction. (**A**-**B**) 10-14 wk old C57BLKS/J db/+ or db/db mice (n=4 mice per genotype) were fasted for 4 h and then injected i.p. with glucose (1 mg/g bw). Blood glucose was monitored for 2 h as indicated (**A**) and plasma sampled for insulin (**B**). (**C**-**H)** Islets isolated from 10-14 wk old C57BLKS/J db/+ vs. db/db were examined for ultrastructure by transmission electron microscopy (n=3 mice per genotype). β-cells were identified morphologically by the presence of insulin granules. (**C**) Representative micrographs highlighting Golgi stacks (red arrows) are shown. Images were quantified for cisternal length (**D**), cisterna number per stack (**E**), Golgi stacks per cell (**F**), swollen cisternae (**G**), and degree of vesiculation (**H**). Number of images analyzed per genotype is listed. (**A, B, D**-**H**) Data represent the mean ± S.E.M. * p < 0.05 by two-way ANOVA, repeated measures (**A**-**B**) or Student t test (**D**-**H**).

## Discussion

A fundamental feature of the β-cell’s secretory capacity is the ability to match the demands of insulin release with the production of insulin granules^2,8^. Although the β-cell may only secrete 5-10 % of the total insulin content per hour^3,4^, the strong preference for release of newly synthesized insulin highlights the importance of defining critical determinants in the proinsulin biosynthetic pathway^5^. Much of our current understanding of the cellular mechanisms regulating secretory protein biogenesis stems from early work utilizing pulse-chase radioisotope labeling of prohormones, such as proinsulin and prolactin, combined with ultrastructural analysis to define the stepwise transition between secretory organelles^38^. These studies identified key regulatory steps during proinsulin transit through the secretory pathway (ER, Golgi, granule) and defined the maturing insulin granule as the primary site for proinsulin to insulin conversion^14,26^. Extending from the concept of pulse-chase labeling, fluorescent reporter systems, such as SNAPtag and RUSH, have recently been developed to further evaluate protein trafficking dynamics using light microscopy. For example, RUSH-based reporters have demonstrated that the endolysosomal proteins, mannose-6-phosphate receptor and LAMP1, are segregated early in the Golgi and bud from distinct TGN domains into separate transport carriers^39^. Furthermore, SNAPtag labeling of proinsulin has revealed age-dependent differences in granule mobility that may explain the high probability for utilizing nascent insulin granules for exocytosis^27^. In the current study, we have used a proinsulin-based RUSH reporter, proCpepRUSH, to evaluate proinsulin transit through the secretory system of primary β-cells. Using the synchronized release of proCpepRUSH from the ER, we show that the kinetics of proCpepRUSH trafficking from the ER to secretory vesicles occurs within 2 h of biotin addition. During proCpepRUSH transit through the Golgi, we observed proCpepRUSH condensates, which are consistent with proinsulin segregation occurring early in Golgi maturation^14,40^. We further demonstrate that processing of proCpepRUSH to CpepRUSH occurs in post-Golgi vesicles, and that only fully processed CpepRUSH can be secreted from β-cells in a glucose-dependent manner. Collectively, the proCpepRUSH reporter recapitulates the essential steps of proinsulin trafficking and maturation in primary β-cells and is a valuable tool for deciphering insulin granule dynamics in normal and disease states.

Defects in β-cell function directly contribute to the development of hyperglycemia in major forms of diabetes due to the singularity of insulin’s role as the only hormone capable of reducing blood glucose^23,36,41^. Because insulin granule production is a defining feature of β-cell secretory capacity, alterations in proinsulin biosynthesis may be a critical determinant in the etiology of β-cell dysfunction in diabetes^2,8,37,42^. Using a rodent dietary model of β-cell dysfunction, we recently demonstrated that a delay in proinsulin trafficking from the ER contributes to insulin insufficiency by limiting available proinsulin for insulin granule formation^10,12^. In the current work, we report that insulin granule trafficking from the Golgi is also impaired in a rodent diabetes model and coincides with morphological alterations to Golgi structure. Using a temperature block to synchronize proinsulin exit from the TGN, we show that insulin-containing granules remain proximal to the Golgi, which may be the result of a Golgi exit delay. Indeed, this trafficking defect is consistent with the appearance of swollen Golgi cisternae, which may harbor proinsulin cargo that are delayed in Golgi exit. Additionally, the reduced appearance of newly-synthesized proCpepRUSH positive vesicles at the plasma membrane is consistent with a past report identifying delays in plasma membrane docking of nascent granules in human T2D β-cells^43^. Whether the decreased docking of nascent insulin granules is a direct result of delayed Golgi exit, or represents a separate defect in post-Golgi trafficking remains to be determined.

The morphological change in Golgi structure contrasts with the observation that ER structure appears relatively normal in β-cells from human T2D and rodent diabetes models, including db/db mice used in this study^8,12,44–46^. Despite the strong association of ER stress with the onset of β-cell dysfunction^47^, morphological changes to ER structure are relatively modest in T2D and largely include expansion of the ER membrane surface, rather than dilation of the ER lumen^12,45,46^. ER redox imbalance and proinsulin misfolding in T2D can lead to the formation of disulfide-linked aggregates in the ER^10–12^. Presumably, ER-associated degradation (ERAD) and ER autophagy are utilized to eliminate some aggregates^48–50^; however, a portion of misfolded proinsulin may exit the ER and accumulate in the Golgi leading to the swollen cisternae observed here. In support of this, lysosomal degradation of post-Golgi vesicles containing proinsulin has been observed in human and rodent T2D β-cells and may represent a clearance pathway for removal of misfolded proinsulin residing in defective insulin granules^51^. While the molecular mediators of ER stress are well-documented^47,52^, very little is known about how Golgi stress manifests, and if there is a distinct transcriptional program like the ER stress response that actively serves to restore Golgi functions^2,53,54^.

Compromised Golgi integrity due to loss of the ribbon structure and/or fragmentation of the Golgi stacks has been documented in other protein misfolding diseases, such as Alzheimer’s disease^55–58^. These alterations can lead to protein mis-sorting and export defects as well as alterations to post-translational modifications such as lipidation and glycosylation. Protein mis-sorting in the development of T2D has long been suggested based on the observation that inappropriate release of proinsulin from the β-cell, i.e. hyperproinsulinemia, is a common clinical hallmark^37,59,60^. In addition, individuals with established T1D present with elevated circulating proinsulin compared to C-peptide levels^23,61^, suggesting that dysregulation of secretory protein trafficking and maturation occurs in the few remaining β-cells in autoimmune diabetes as well^62^. While changes in Golgi structure have not been reported in T1D, the mis-trafficking of proinsulin with cathepsin D and formation of chromogranin A-insulin hybrid peptides illustrates that Golgi sorting defects could be a central underlying theme of β-cell pathophysiology^63,64^. Future studies will continue to explore the contribution of Golgi function to proinsulin trafficking and whether defects in Golgi protein sorting and export contribute to impaired insulin secretion in both major forms of diabetes.

## Experimental methods

### Cell culture and Reagents

832/3 cells (a gift from Dr. Christopher Newgard) were cultured as previously described (cite). For introduction of adenoviral reporters, cells were transduced with ∼1-5 × 10^7^ IFU/mL adenovirus for 18 h and assayed 72-96 h post-treatment. Cell culture reagents were from Thermo Life Technologies unless specified otherwise. Chemical reagents were from Sigma-Aldrich unless specified otherwise. Mouse islets were isolated via collagenase V digestion and purified using Histopaque 1077 and 1119. Islets were cultured in RPMI supplemented with 10% fetal bovine serum and 1% penicillin and streptomycin and maintained at 37°C in 5% CO_2_. Pools of islets were transduced with ∼ 1-5 × 10^8^ IFU/mL adenovirus for 18 h and assayed 72-96 h post-treatment.

### Animal Studies

Male C57BL6/J mice (8-10 wk old: Jackson Laboratories) were placed on Western diet (Research Diets, D12079B) or standard chow for 8-10 weeks. Body weight was measured weekly. BKS.Cg-Dock7^m^ (C57BLKS/J) Lepr^db/+^ and Lepr^db/db^ mice (db/+, db/db) were generated by heterozygous cross and genotypes confirmed via real time PCR according to Jackson Laboratories. Hyperglycemic db/db mice (*ad* lib fed blood glucose > 220 mg/dl) were compared to age-matched (10-14 weeks old), normoglycemic littermate controls (db/+). Glucose tolerance was measured in 4-6 h fasted mice given a 1 mg/g body weight (i.p.) challenge. Blood glucose was determined using a One Touch Ultra 2 glucometer. Plasma insulin was determined by ELISA (ALPCO 80-INSMSU-E10). All animal protocols were approved by the University of Iowa Institutional Animal Use and Care Committee.

### Plasmids and Viruses

ProCpepRUSH (SA-KDEL-IRES-proCpep-SBP-GFP) was generated by Gibson assembly and subcloned into pENTR2b-RIP^12^. SA-KDEL-IRES and SBP PCR fragments were separately amplified from Str-KDEL_SBP-EGFP-GPI (gift from Franck Perez, Addgene plasmid #65294; RRID:Addgene_65294)^28^. sfGFP was PCR amplified from sfGFP-C1 (gift from Michael Davidson and Geoffrey Waldo, Addgene plasmid #54579; RRID:Addgene_54579)^65^. Proinsulin was PCR amplified as two fragments on either side of the ApaI site from human *INS* cDNA for insertion of the SBP and sfGFP fragments within the C-peptide coding sequence. RIP-SA-KDEL-IRES-proCpep-SBP-GFP was recombined from pENTR2b-RIP into a modified pAD-PL/DEST via Gateway cloning using LR Clonase II^66^. Recombinant adenoviruses were generated in HEK293 cells and purified by cesium chloride gradient. All sequences were verified by the Iowa Institute of Human Genetics, University of Iowa.

### Glucose-stimulated insulin secretion

Static incubation for insulin secretion was performed on pools of 10 islets in secretion assay buffer^67^ containing 2.5 mM glucose for 1 h at 37° C followed by incubation at 12 mM glucose for 1 h. Insulin was measured by ELISA (rodent 80-INSMR-CH10; ALPCO). For determination of proCpepRUSH secretion, INS1 832/3 cells expressing proCpepRUSH were incubated in secretion assay buffer without BSA, containing 2.5 mM glucose for 3 h at 37° C or 12 mM glucose for 3 h. Media was precipitated by addition of ice-cold 10 % (v/v) trichloroacetic acid (TCA) and examined by immunoblot as described below. Cells were lysed in RIPA buffer and total protein determined using BCA (Thermo Life Technologies).

### Immunoblot Analysis

Clarified cell lysates and TCA precipitated media were resuspended in LDS sample buffer (Thermo Life Technologies), resolved on 4-12 % NuPAGE gels (Thermo Life Technologies) and transferred to supported nitrocellulose membranes (BioRad). Membranes were probed with diluted antibodies raised against GFP (rabbit, Abcam ab290) or γ-tubulin (mouse, Sigma, T5326). Donkey anti-mouse and anti-rabbit antibodies coupled to IR-dye 680 or 800 (LI-COR) was used to detect primary antibodies. Blots were developed using an Odyssey CLx Licor Instrument. Original uncropped blots are provided as a supplemental data file.

### Quantitative RT-PCR

RNA from mouse islets were harvested using the RNeasy Microkit (QIAGEN) and cDNA synthesized using iScript (BioRad) or Lunascript (NEB). Real-time PCR was performed using the QuantStudio-7 PRO sequence detection system and software Design & Analysis. All primer sequences are provided as a supplemental data file.

### Fluorescence Microscopy and Imaging

Isolated islets expressing proCpepRUSH (AdRIP) were gently dispersed into monolayer sheets using Accutase (Sigma-Aldrich) and plated onto HTB9 coated coverslips, 6 cm glass bottom dishes (Mattek) or 24-well plates as previously described^18,68^. INS1 832/3 cells expressing proCpepRUSH were plated on HTB9 coated coverslips at low density and cultured overnight. To initiate proCpepRUSH trafficking, biotin was added to a final concentration of 200 µM in the culture media and fixed in 10 % neutral-buffered formalin at the indicated times (30-180 min). In specified experiments, islet cells were cultured at 19° C for 1 h. Biotin was added to the culture media (200 µM) and cells incubated at 19° C for an additional 2 h. Cells were then shifted to 37° C for 2-3 h as indicated and fixed in 10 % neutral-buffered formalin. For immunostaining, permeabilized cells were incubated overnight with antibodies raised against cation independent-M6PR (rabbit, Proteintech 20253-1-AP), CgB (rabbit, Proteintech, 14968-1-AP), CPE (rabbit, Proteintech 13710-1-AP), GM130 (mouse, BD Transduction 610823), GRP94 (rabbit, kind gift of Dr. Christopher Nicchita, Duke University), insulin (guinea pig, Fitzgerald, 70R-10659), SA (rabbit, Rockland 100-4195), and TGN38 (mouse, Novus Biologicals NB300-575) as indicated. Highly cross-adsorbed fluorescent conjugated secondary antibodies (whole IgG, donkey anti-guinea pig AlexaFluor488, donkey anti-rabbit Cy3, and donkey anti-mouse AlexaFluor 647; Jackson ImmunoResearch) were used for detection. Cells were counterstained with DAPI and mounted using Fluorosave (Calbiochem).

For granule distance measures, images were captured on a Leica SP8 confocal microscope using a HC PL APO CS2 63x/1.40 oil objective with 3x zoom as z-stacks (5 per set, 0.3 μm step, 0.88 μm optical section) and deconvolved (Huygen’s Professional). Granule distance measurements from the Golgi were determined using a distance transformation module in Imaris (Bitplane) from spot-rendered proCpepRUSH-positive granules (150-300 nm) and surface rendering of the Golgi identified through GM130 immunostaining. Granule distances were binned as indicated and expressed as a percentage of the total to normalize between cells and conditions ^12,19^.

For plasma membrane detection, proCpepRUSH-labeled immunostained (fixed) cells were maintained in PBS and imaged using a Leica TIRF AM microscope via a 100× oil objective in TIRF mode with a penetration depth of 150 nm. ProCpepRUSH-positive granule numbers and proCpepRUSH-expressing cells (defined by nuclei) were determined using Fiji/ImageJ. SNAP25 immunostaining was used to identify the plasma membrane.

For time lapse images, mouse islets expressing proCpepRUSH (AdRIP) were plated on HTB9-coated 6 cm glass bottom dishes (Mattek). Images were acquired every 30 s using a Zeiss 980 AiryScan2 confocal microscope with heated stage and lid (37°C) and 5 % CO_2_ using a 40x water objective (NA=1.2) objective. Images were collected as z-stacks (7 slices per stack, d=1 µm) and processed using Zen (Zeiss) and Imaris (Bitplane) software.

### Ultrastructure

All EM related reagents were from Electron Microscopy Sciences (EMS; Hatfield, PA). Isolated islets were PBS washed and fixed in 2.5 % glutaraldehyde, 4 % formaldehyde cacodylate buffer overnight (16-18 h) at 4° C. Tissue was post-fixed in fresh 1 % OsO_4_ for 1 h, dehydrated using a graded alcohol series followed by propylene oxide and embedded in Epon resin as previously described ^55^. Resin blocks were trimmed with a glass knife, cut to ultrathin (50-70 nm) sections with a diamond knife, and mounted on Formvar-coated copper grids. Grids were double contrasted with 2 % uranyl acetate then with lead citrate. Images were captured at 1,500x, 3,000x, 6,000x, and 8,000x magnifications by a JEOL JEM-1400 transmission electron microscope.

### Statistical Analysis

Data are presented as the mean ± S.E.M. For statistical significance determinations, data were analyzed by the two-tailed unpaired, Student’s t test or by ANOVA with post-hoc analysis for multiple group comparisons as indicated (GraphPad Prism). A *p*-value < 0.05 was considered significant.

## Supporting information

Time lapse recording of proCpepRUSH trafficking

## Author contributions

C.K.B., and S.B.S conceived and designed studies. C.K.B., C.J.B. and S.B.S collected samples, performed the experiments, and analyzed the data. J.Z., and Y.W. performed TEM. C.K.B. and S.B.S. wrote the manuscript.

## Acknowledgements

We would like to thank Kristen Rohli and Sandra Blom for helpful comments and discussion and Thomas Moninger for expert technical assistance. We would like to acknowledge the use of the University of Iowa Central Microscopy Research Facility. This work was supported by National Institutes of Health grant R35GM130331 to Y.W. and Department of Defense Congressionally Directed Medical Research Program grant W81XWH-20-1-0200 to S.B.S. S.B.S. is the guarantor of this work and, as such, had full access to all the data in the study and takes responsibility for the integrity of the data and the accuracy of the data analysis. All data are available upon reasonable request.

## Abbreviations

(ER): endoplasmic reticulum
(Glc): glucose
(GSIS): glucose-stimulated insulin secretion
(IFU): infectious units
(IRES): internal ribosome entry site
(RIP): rat insulin promoter
(RUSH): retention using selective hooks
(SC): standard chow
(SA): streptavidin
(SBP): streptavidin-binding peptide
(sfGFP): superfolder GFP
(TGN): transmission electron microscopy
(TEM): *trans-*Golgi network
(TCA): trichloroacetic acid
(T2D): Type 2 diabetes
(WD): western diet
(TIRF): total internal reflection fluorescence

## Supplemental figure legends

**Supplemental Figure 1.**
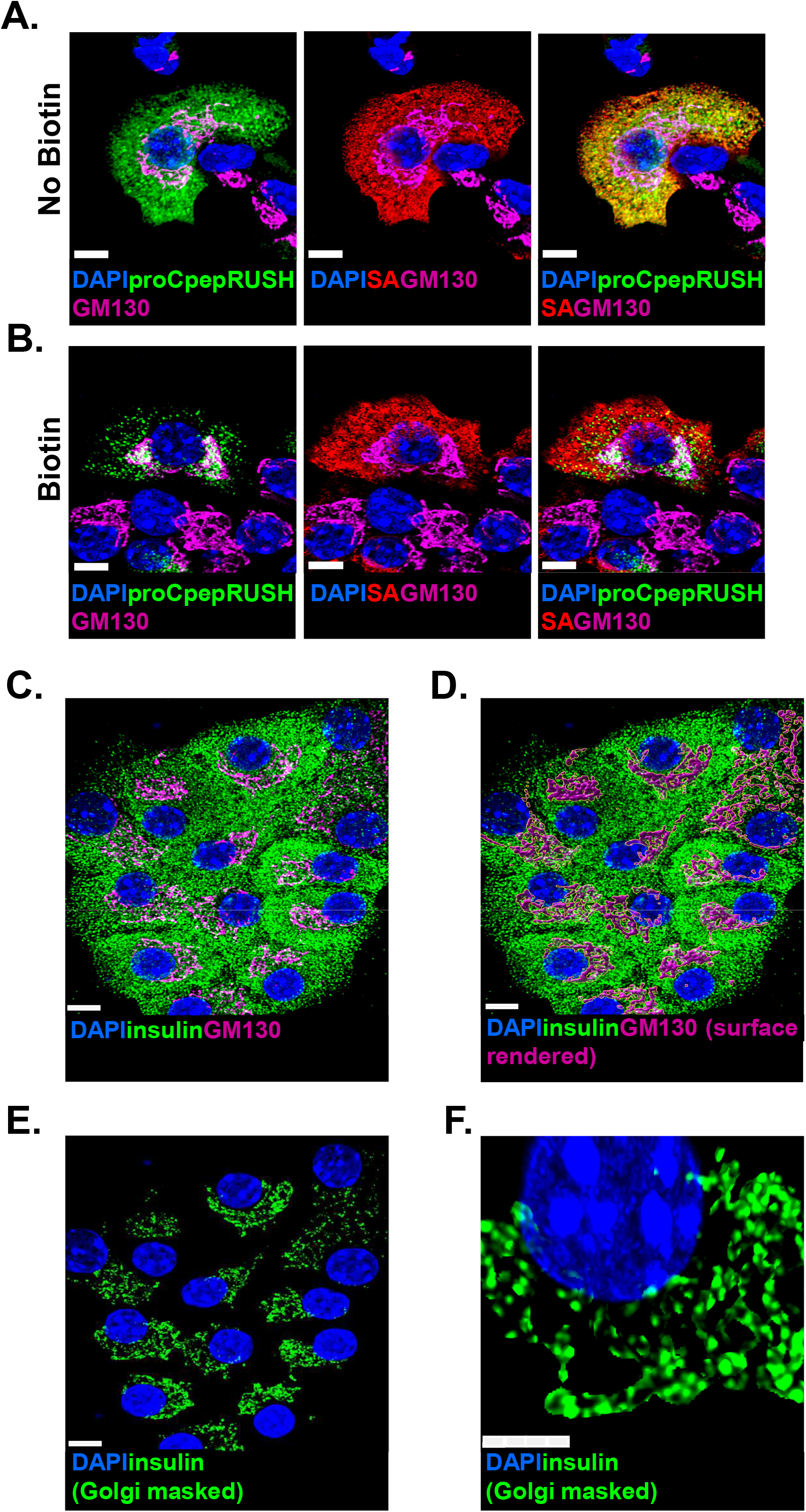
Proinsulin puncta condense in the Golgi. (**A**-**B**) Mouse islets (C57BL6/J) treated with AdRIP-proCpepRUSH were examined 48 h post-infection. proCpepRUSH (green) localization was examined before (no biotin) or after biotin treatment (200 μM, > 3h). Cells were immunostained for SA-KDEL (red), GM130 (magenta) and counterstained with DAPI (blue). (**C**-**F**) Isolated mouse islets were immunostained for insulin (pan-insulin antibody; green), TGN38 (magenta) and counterstained with DAPI (blue). (**C**) Representative image is shown for immunostaining. 3D-surface rendering of the GM130 volume was used as a mask (**D**) for image segmentation to specifically examine proinsulin staining within the Golgi (**E**). (**F**) Additional magnification used to highlight proinsulin staining within the Golgi. Scale bar = 5 μm. Related to Figure 1.

**Supplemental Figure 2.**
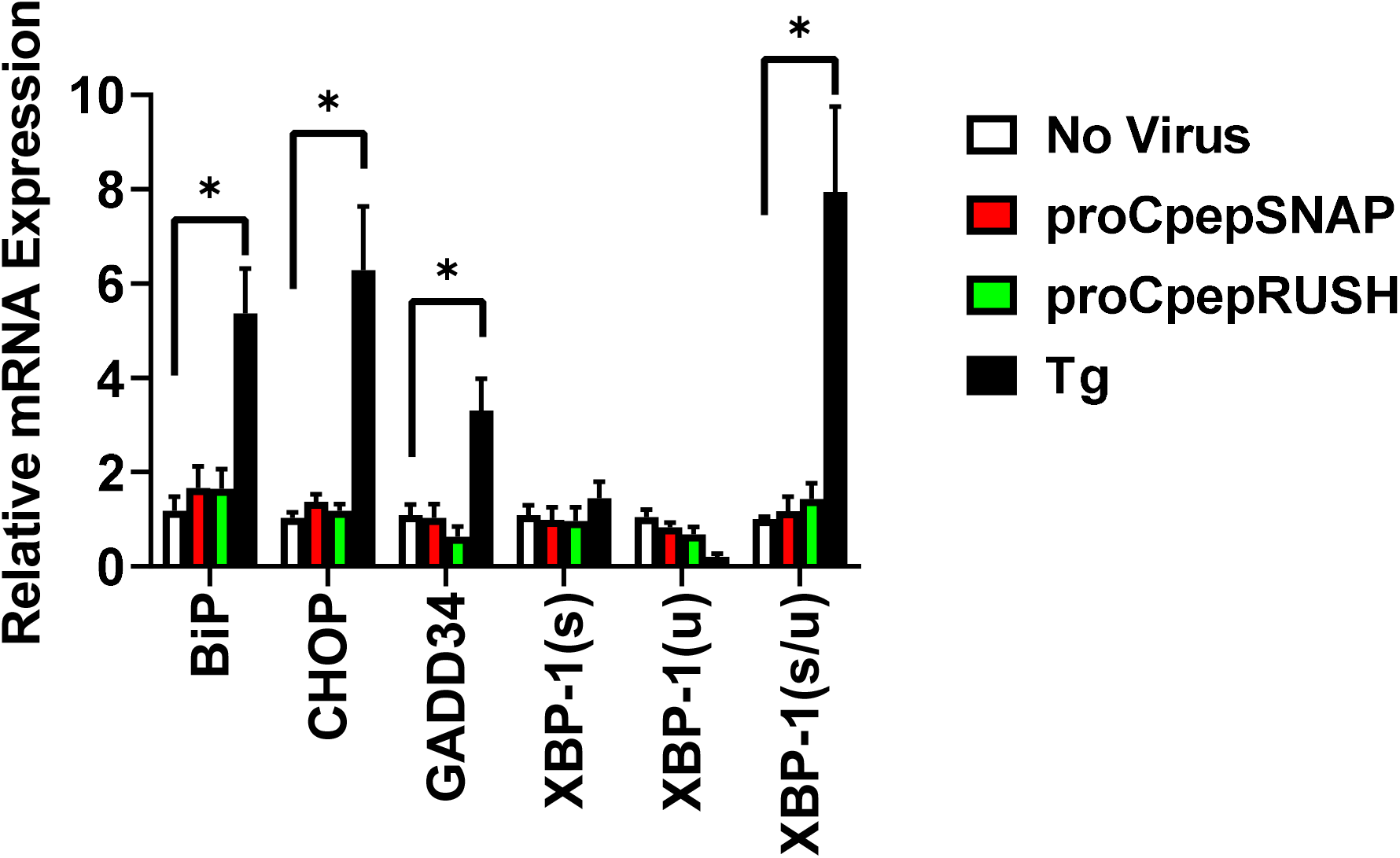
proCpepRUSH does not elicit the ER stress response. Isolated mouse islets were treated with AdRIP-proCpepRUSH, AdRIP-proCpepSNAP or no virus. 72 h post-infection, mRNA expression was examined and compared to islets treated for 18 h with thapsigargin (500 nM) as indicated. Data represent the mean ± S.E.M. *p < 0.05 by 2 way-ANOVA with Sidak post-test analysis. Related to Figure 3.

**Movie S1. Time-lapse video of proCpepRUSH trafficking in primary β-cells**. Mouse islets (C57BL6/J) treated with AdRIP-proCpepRUSH were examined 48 h post-infection. Biotin addition (200 μM) was used to initiate proCpepRUSH (green) trafficking. Image acquisition begins 12 min post-biotin treatment. Scale bar = 5 µm. Related to Figure 1.

